# Food pairing independently associated with cardiometabolic traits beyond dietary patterns across western and eastern populations

**DOI:** 10.1101/2025.01.03.631167

**Authors:** Qiufeng Deng, Yuhao Sun, Mingxia Gu, Jingxiang Fu, Fengzhe Xu, Yuwen Jiao, Yan Zhou, Beining Ma, Lu Liu, Xiuchao Wang, Quanbin Dong, Tingting Wu, Huayiyang Zou, Jing Shi, Yifeng Wang, Yanhui Sheng, Liming Tang, Wei Sun, An Li, Shixian Hu, Jusheng Zheng, Yan He, Hongwei Zhou, Wei Wu, Xiangqing Kong, Lianmin Chen

**Affiliations:** Department of Cardiology, The First Affiliated Hospital of Nanjing Medical University, Nanjing Medical University, Nanjing, China; Changzhou Medical Center, The Affiliated Changzhou No.2 People’s Hospital of Nanjing Medical University, Nanjing Medical University, Changzhou, China; Cardiovascular Research Center, The Affiliated Suzhou Hospital of Nanjing Medical University, Suzhou Municipal Hospital, Gusu School, Nanjing Medical University, Suzhou, China; Microbiome Medicine Center, Department of Laboratory Medicine, Zhujiang Hospital, Southern Medical University, Guangzhou, China; School of Life Sciences, Westlake University, Hangzhou, China; Department of Periodontology, Stomatological Hospital, School of Stomatology, Southern Medical University, Guangzhou, China; Institute of Precision Medicine, The First Affiliated Hospital, Sun Yat-sen University, Guangzhou, China; Guangdong Provincial Institute of Public Health, Guangdong Provincial Center for Disease Controland Prevention, Guangzhou, Guangdong, China

**Keywords:** population study, dietary foods, long-term food pairing patterns, gut microbiome, cardiometabolic health

## Abstract

Cardiometabolic health is closely linked to diet. Both specific food choices and dietary indices are commonly used to guide cardiometabolic protection strategies. However, the balance between foods may also significantly impact cardiometabolic health.

Here, we explore the balance and imbalance in long-term dietary food intake to quantify food pairing patterns (FPs), providing insights beyond conventional dietary indices and single-food intake frequencies. Using data from our GGMP (n = 6,994) and the NHANES (n = 7,350) cohorts, we observed that long-term food pairing patterns are independent of single food intake frequency and dietary indices. We identified 1,759 and 306 cardiometabolic-related long-term food pairing patterns from two cohorts (FDR < 0.05), respectively. Notably, around 80% of these pairing foods not individually associated with cardiometabolic traits, and food pairing pattern associations with cardiometabolic traits at hyper food group level were consistent across Eastern and Western populations. Besides, mediation analysis revealed that 72.7% of long-term food pairing patterns affected cardiometabolic traits through 31 microbial genera, with *Clostridiumsensustricto1* playing a predominant role. Moreover, multi-step mediation analysis found that microbes mediated the impact of long-term food pairing patterns on cardiometabolic traits primarily through their metabolic pathways, such as pyruvate fermentation to propanoate and ergothioneine biosynthesis pathways.

Our data suggest that balance between dietary foods is broadly associated with cardiometabolic traits by modulating gut microbial functionalities, in contrast to the lower occurrence when considering individual foods in isolation. These results offer a novel perspective for designing personalized dietary strategies beyond present dietary indices to enhance cardiometabolic health.

## 1. Introduction

Nutrition is a well-established influencing factor for human cardiometabolic health[1,2]. It has been widely accepted that both the intake of specific foods and dietary pattern indices are associated with cardiometabolic health. For instance, cohort-based studies have shown that the intake of specific foods, such as red meat, can increase cardiovascular mortality risk[3–5], while the DASH (Dietary Approaches to Stop Hypertension) dietary pattern index[6] and MED (Mediterranean Diet) index[7] are negatively associated with cardiovascular mortality risk. It is important to note that the former confirms the importance of specific foods in disease prevention, while the latter illustrates the cumulative effect of a group of foods.

Nevertheless, in addition to those two aspects, the daily diet of each individual encompasses a diverse array of foods that exhibit clear regional differences. For instance, Western settings exhibit a predominantly Western diet characterized by a higher intake of processed meat, red meat, butter, high-fat dairy products, and refined grains[8,9]. In contrast, Eastern settings are characterized by an Eastern diet, with higher intakes of whole grains, vegetables, and seafood[10]. Importantly, different foods are not metabolized independently but undergo complicated interactions that are still far from fully understood[11]. The balance and imbalance of different food intakes can regulate the composition of the gut microbiome and host metabolism, ultimately influencing human health. For example, maintaining a balance between vitamin-enriched fruits and fiber-enriched vegetables has been shown to reduce the risk of all-cause mortality[12]. In different food combinations, the characteristic gut microbiome and its metabolites may coordinate or mediate the occurrence and development of cardiovascular diseases, obesity, and other chronic diseases[13–17]. Thus, investigating food pairing patterns and their associations with individual health and disease may enhance our understanding of personalized nutrition from an ecological perspective. However, investigation of food pairing patterns and their impact on human cardiometabolic health remains limited.

In this study, we aim to elucidate the effects of balanced and imbalanced long-term food pairing patterns on human cardiometabolic phenotypes between populations with different dietary preferences (**Figure 1**). Additionally, we further interpret how long-term food pairing patterns influence cardiometabolic health from the perspective of the gut microbiome and its metabolic functionalities. This study offers a unique angle on understanding the role of diet and ultimately assist in designing personalized diet to improve human health.

**Figure 1.**
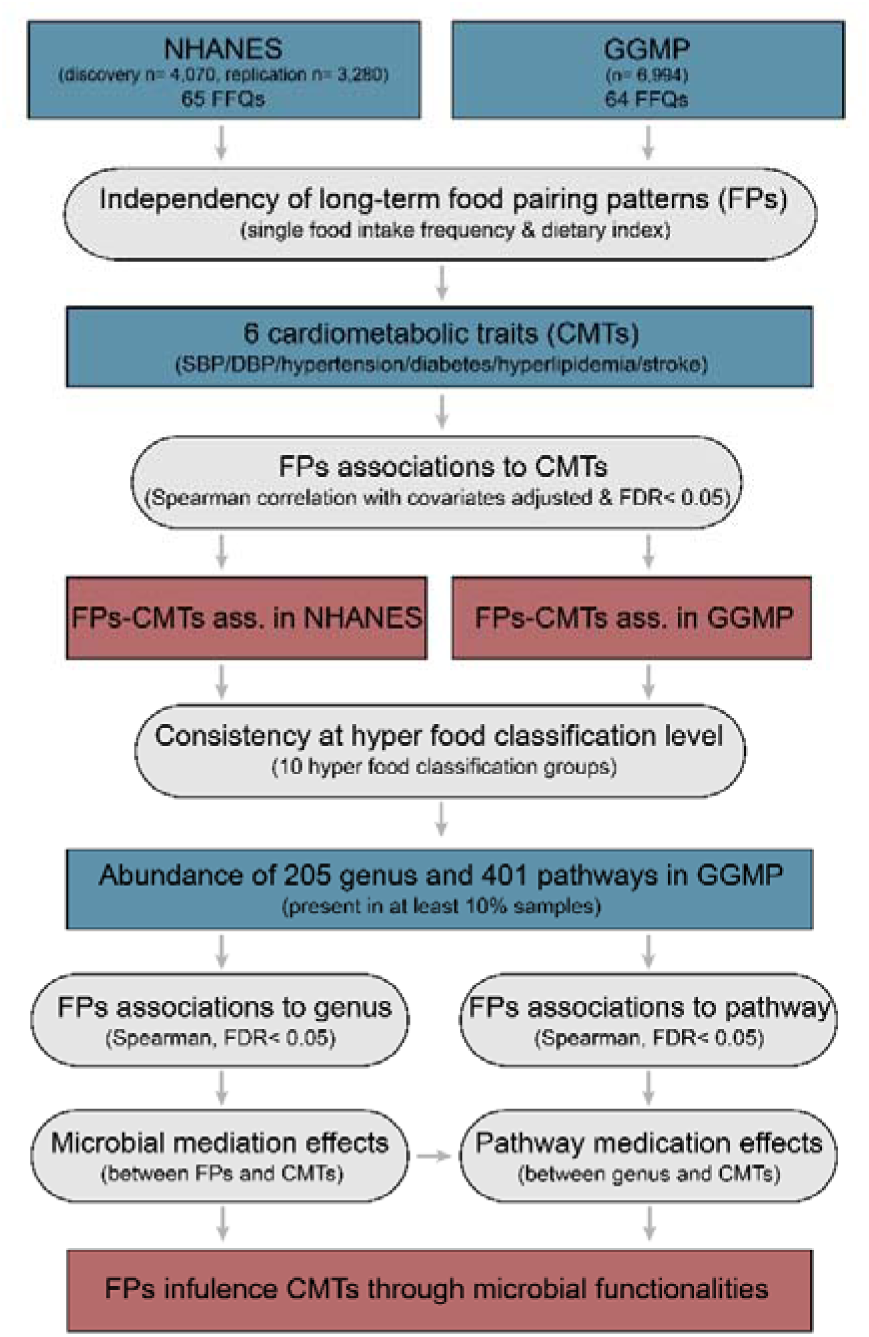
Overview of the study design and analysis workflow. This study involves two large-scale prospective cohorts, including the NHANES (National Health and Nutrition Examination Survey, n = 7,350) and the GGMP (Guangdong Gut Microbiome Project, n = 6,994). A total of 65 and 64 normalized unique monthly food intake frequencies were collected based on food frequency questionnaires (FFQ) from the NHANES and GGMP cohorts, respectively. Additionally, 6 cardiometabolic traits (CMTs) were assessed, including systolic blood pressure (SBP), diastolic blood pressure (DBP), and the prevalence of hypertension, diabetes, hyperlipidemia, and stroke. For the GGMP cohort, gut microbiome data were also available.

## 2. Methods

### 2.1. Studying cohorts

In this study, two large scale population-based cohorts have been involved, including the NHANES (National Health and Nutrition Examination Survey, n = 7,350) and the GGMP (Guangdong Gut Microbiome Project, n = 6,994). NHANES was conducted by the National Center for Health Statistics (NCHS) to obtain data on the health and nutritional status of the non-institutionalized U.S. population. NHANES used a complex, multi-stage, stratified, clustered sampling design to achieve a nationally representative sample. This nationwide survey included demographic and basic health data, standardized medical examinations, and laboratory tests. The NCHS Research Ethics Review Board approved all the data collection procedures. Detailed descriptions of NHANES are available elsewhere[18]. For this study, we involved 7,350 out of 62,160 NHANES participants from 1999 through 2010 in the United States for whom detailed dietary habit is available (**Table S1**), and 54,810 were not eligible because of missing data on complete food frequency questionnaires. We further separated them into discovery set (n = 4,070; NHANES data from 2005 to 2010) and replication set (n = 3,280; NHANES data from 1999 to 2004) to verify the reproducibility of long-term food pairing patterns in western population. The GGMP cohort (n = 7,009) is a large community-based cross-sectional cohort conducted between 2015 and 2016 from Guangdong Province, China. Detailed information regarding the host metadata and sample collection have been reported previously[19]. The GGMP cohort was 55% female and 45% male, with a mean age of 53 years (s.d. = 15). The mean BMI was 23.4 (s.d. = 3.5) and 26% current smokers. For this study, we involved 6,994 out of 7,009 GGMP participants for whom socio-demographic features, disease status, detailed dietary habit and stool microbiome information is available (**Table S2**). The GGMP cohort study was approved by the Ethical Review Committee of the Chinese Center for Disease Control and Prevention under Approval Notice No. 201519-A. Written consent was also obtained from all participants.

### 2.2. Dietary information from food frequency-based questionnaires

The participants from NHANES completed 65-item food frequency questionnaires (FFQ) to assess the frequencies of specified food items they had consumed over the past time without considering the portion size. 65 food items were classified into ten food categories: 14 grains, 7 meats, 11 milks, 1 egg, 10 vegetables, 4 fruits, 10 drinks, 2 legume, 3 oils, 3 snacks **(Table S3**). The participants from GGMP completed a 64-item FFQ to assess the frequencies of specified food items they had consumed over the past time without considering the portion size. 64 food items were classified into nine food categories: 8 grains, 12 meats, 7 milks, 3 eggs, 8 vegetables, 3 fruits, 11 drinks, 6 legumes, 6 snacks (**Table S4**).

Both FFQs of NHANES and GGMP included common foods, such as rice, beer, coffee, beef, yogurt, wine. However, due to the differences between western and eastern dietary habits, there were also many unique foods in both questionnaires. For example, NHANES cohort contained hot cocoa, margarine, dark bread, etc., and GGMP cohort contained wood fungus, crab, and thousand egg, etc. In the data processing of FFQs for both cohorts, we all normalized the intake frequency to convert the consumption frequency of daily, weekly, quarterly, and annual consumption into monthly consumption frequency for food-related analysis in context.

### 2.3. Construction of long-term food pairing patterns

We hypothesized that the balance and imbalance of food intake, as captured through long-term food frequency questionnaires, could provide unique insights into the role of diet in cardiometabolic health. These aspects are not adequately reflected in traditional single-food intake frequency analyses or dietary indices. Specifically, we proposed two types of food pairing patterns: Additive Food Pairing Pattern (AFP), which assumes synergistic effects between two foods, and Subtractive Food Pairing Pattern (SFP), which assumes antagonistic effects between two foods. The AFP and SFP were calculated using the normalized monthly consumption frequency of individual foods according to the following formulas:

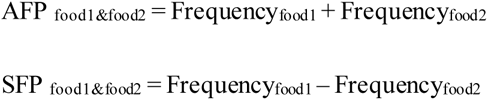

where Frequency_food1_ and Frequency_food2_ denoted the monthly intake frequencies of the two foods being paired. Specifically, AFP reflects the combined consumption frequency of two foods, with higher AFP values indicating more frequent joint consumption. This suggests that the combined dietary impact of the two foods may be amplified. SFP, on the other hand, captures the difference in consumption frequencies between two foods. Positive or negative SFP values indicate imbalances in relative consumption, which may influence dietary effects in distinct ways. Balanced food pairings suggest that the combined consumption frequency of two foods significantly impacts phenotypes, whereas imbalanced pairings indicate that large differences in consumption frequencies between the two foods may uniquely affect phenotypic outcomes.

As we have 65 and 64 normalized food intake frequency traits for NHANES and GGMP, respectively. Based on the above formulas, we finally generated 2080 and 2016 long-term food pairing patterns (AFP and SFP) for the two cohorts separately, thus providing a comprehensive assessment of potential long-term food pairing patterns across cohorts.

### 2.4. Independency of long-term food pairing patterns on top of previous dietary indicators

To investigate the independency of long-term food pairing pattern proposed here, we applied Spearman correlation to assess the correlation between 2080 AFPs/SFPs and other dietary indices, including Healthy Eating Index-2020 (HEI2020)[20], Healthy Eating Index 2015 (HEI2015)[21], Alternative Healthy Eating Index (AHEI)[22], Dietary Approaches to Stop Hypertension (DASH)[23], DASH Index in serving sizes from the DASH trial (DASHI)[24], Alternate Mediterranean Diet Score (aMED)[25], MED Index in serving sizes from the PREDIMED trial (MEDI)[26] and Dietary Inflammation Index (DII)[27].

### 2.5. Long-term food pairing pattern associations to cardiometabolic traits

To assess associations between long-term food pairing patterns and cardiometabolic traits in NHANES cohort, continuous AFPs and SFPs were inverse-rank transformed and corrected for age, sex, race, BMI, and smoking status. Spearman correlation was then applied to assess the correlation between 2080 AFPs/SFPs and 6 cardiometabolic traits (SBP, DBP, hypertension, diabetes, hyperlipidemia and stroke) in the NHANES discovery and replication sets, respectively (**Table S5**). In the GGMP cohort, 2016 AFPs/SFPs were associated with 6 cardiometabolic traits after adjusting for age, sex, BMI and smoking status by using Spearman correlation (**Table S6**). The FDR was calculated using the Benjamini– Hochberg procedure[28].

To further assess the robustness of our findings, a sensitivity analysis was conducted by including diet diversity as an additional covariate. Diet diversity was defined as the total number of different foods consumed per individual and was calculated based on the FFQ data.

### 2.6. Gut microbiome data processing

For the GGMP cohort study, participants received a stool sampler, ice bag, and instructions for proper sample collection and storage. After defecation, participants placed the fecal sample with the ice bag in the provided ice box and sent it to a nearby collection point within 24 hours. Collected samples were further transferred to the laboratory and stored in –80 °C freezers until further processing. Total bacterial DNA was extracted using a Fecal DNA Bead Isolation kit (Bioeasy, Shenzhen) according to the manufacturer’s instructions and the 16 S rRNA gene V4 region was amplified and sequenced on an Illumina Hiseq 2500 (Beijing Genome Institute, Beijing, China).

We further performed taxonomic classification using direct taxonomic binning instead of operational taxonomic unit (OTU) clustering methods by taking the advantages of the recent MiBioGen consortium pipeline[29]. In brief, sequence quality-based read filtering was performed using Fastp (v0.23.2). Clean reads were randomly rarefied to 10,000 reads per sample and sequences from all samples were merged into one file. Read identification was performed by using RDP Classifier (v2 .12) based on the SILVA database (v128) and the read labels of each sequence for each sample were summarized. The microbial metabolic pathways were annotated by the PICRUSt2 pipeline[30], which predicts the function of gene sequences and infer the abundance of pathways based on the OTUs from QIIME2 [31]. Finally, 205 genera and 401 pathways with a prevalence of more than 10% were included for downstream analysis.

### 2.7. Microbial associations to long-term food pairing patterns and cardiometabolic traits

We hypothesized a potential connection between the gut microbiome, long-term food pairing pattern, and cardiometabolic health. To assess associations between long-term food pairing patterns and gut microbiome, Spearman correlation was applied to assess the correlation between 2016 AFPs/SFPs and the relative abundance of 205 gut microbial genera in GGMP cohort, adjusted for age, sex, BMI and smoking status (**Table S7**). To further determine the association between specific foods and gut microbiome, we linked dietary frequency to gut microbial genera with Spearman correlation, adjusted for age, sex, BMI and smoking status (**Table S8**). The FDR was calculated using the BH method[28].

Next, to test associations between gut microbiome and 6 cardiometabolic traits, we conducted Spearman correlation analysis to identify the correlation between the relative abundance of 205 gut microbial genera (adjusted for age, sex, BMI and smoking status) and cardiometabolic health (**Table S9**) in GGMP cohort. The FDR was calculated using the BH method.

### 2.8. Mediation analysis

For microbial features associated with both long-term food pairing patterns (FDR < 0.05) and cardiometabolic traits (FDR < 0.05), we first checked whether the long-term food pairing patterns were associated with the 6 cardiometabolic traits using Spearman correlation (FDR < 0.05). Next, we carried out mediation analysis using the mediate function from mediation (version 4.5.0)[32] R package to infer the mediation effect of microbiome for long-term food pairing patterns impacting on cardiometabolic traits (**Table S10**). The FDR was calculated based on the BH method.

### 2.9. Linking the microbial taxa to metabolic pathways

We initially verified the gut microbial genera mediator identified in the above mediation analysis and assessed their associations with gut microbial pathways. Next, we selected microbial metabolic pathways showing statistical significance (FDR < 0.05) to delve deeper into their associations with cardiometabolic traits using Spearman correlation analysis (FDR < 0.05). We then carried out mediation analysis using the mediate function from mediation (version 4.5.0)[32] R package to infer the mediation effect of microbial pathways for mediator genera impacting on cardiometabolic traits (**Table S11**). The FDR was calculated based on the BH method.

## 3. Results

### 3.1. Long-term food pairing patterns show independence from both single food intake frequency and dietary indices

The impacts of diet on cardiometabolic health are primarily focused on the intake of individual foods and dietary patterns reflected by various dietary indices. However, understanding the balance and imbalance of different food intakes may provide additional explanations for cardiometabolic health. Here, we propose long-term food pairing pattern, which employs addition and subtraction to establish configurable connections from one diet to another, including additive food pairing pattern (AFP) and subtractive food pairing pattern (SFP) (see Methods). Specifically, the higher the AFP, the more frequent the intake of either/both diets, and the higher the SFP, the greater the consumption frequency of one food compared to another.

In this manner, we have generated 2080 and 2016 long-term food pairing pattern traits based on 65 and 64 food frequencies from the NHANES (National Health and Nutrition Examination Survey)[18] and our GGMP (The Guangdong Gut Microbiome Project)[33] cohorts, respectively. It is noteworthy that there were only 6 overlapping food frequency questions (rice, yogurt, beef, beer, wine, coffee) between the two cohorts (**Table S3&S4**), suggesting remarkable differences in dietary habits between Eastern and Western populations. By further comparing the distribution of 15 overlapping long-term food pairing patterns between populations, we observed that only the AFP between yogurt and beef intake frequency was comparable between populations (P_Wilcoxon_= 0.08, **Figure 2A**), while all the others were different (**Figure S1**, FDR<0.05). For instance, pairings between rice and beer exhibited significant differences between populations. This was evidenced by a lower total intake of these two foods (as indicated by a lower AFP between rice and beer) and a more balanced consumption frequency (SFP closer to 0) in the US compared to China (**Figure 2A**).

**Figure 2.**
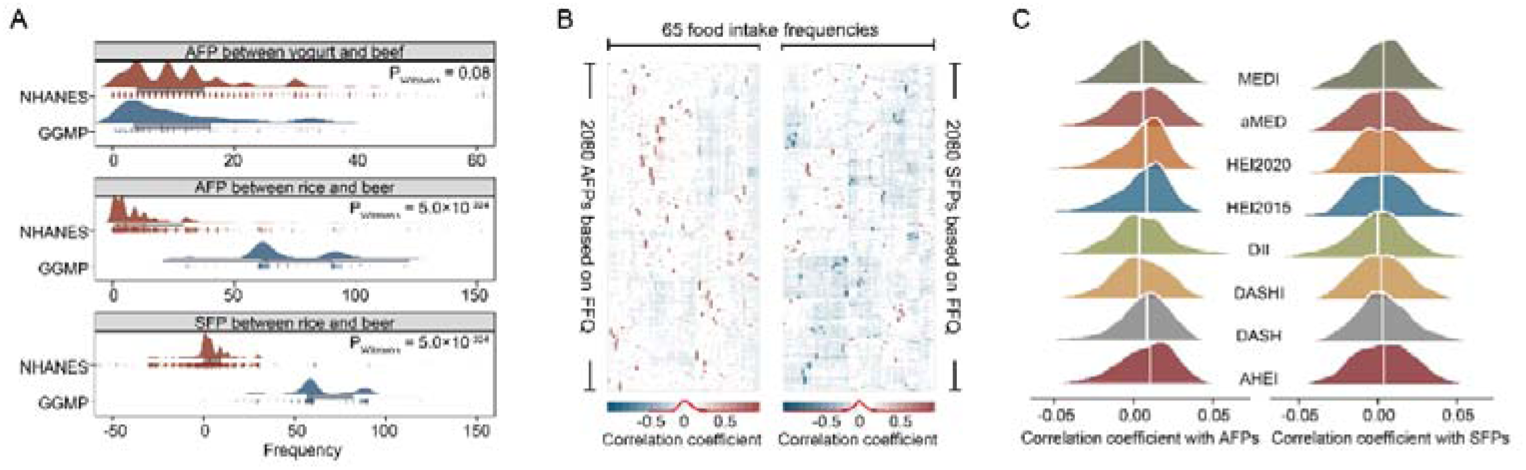
Long-term food pairing patterns independent from both single food intake frequency and dietary pattern. **A.** Distribution of specific long-term food pairing patterns (FPs) across populations, presented as a density map (top), boxplot (middle), and jitter scatter plot (bottom). The boxplot displays the median, first quartile, and third quartile (25th and 75th percentiles) of the FP levels. Whiskers extend to the smallest and largest values within 1.5 times the interquartile range (IQR). Each dot represents one sample. P values from the Wilcoxon test are shown accordingly. **B.** Heatmap illustrating associations between 65 single food intake frequencies and 2080 FPs, estimated using Spearman correlation. The intensity of the color represents the correlation strength. **C.** Distribution of associations between 8 dietary indices and 2080 FPs, estimated using Spearman correlation. FFQ, food frequency questionnaire; MEDI, Mediterranean Diet Index in serving sizes from the PREDIMED trial; aMED, Alternate Mediterranean Diet Score; HEI2020, Healthy Eating Index-2020; HEI2015, Healthy Eating Index-2015; DII, Dietary Inflammation Index; DASH, Dietary Approaches to Stop Hypertension; DASHI, DASH Index in serving sizes from the DASH trial; AHEI, Alternate Healthy Eating Index.

Furthermore, we tested if the long-term food pairing patterns proposed here show independence from other diet-related indicators by assessing the association of food pairing patterns with single diets and dietary pattern indices. 2080 long-term food pairing patterns derived from 65 single food intake frequencies in the NHANES cohort were weakly correlated with both the frequency of a single food intake (**Figure 2B**) and 8 widely recognized diet indices (**Figure 2C**), including Healthy Eating Index-2020 (HEI2020)[20], Healthy Eating Index 2015 (HEI2015)[21], Alternative Healthy Eating Index (AHEI)[22], Dietary Approaches to Stop Hypertension (DASH)[23], DASH Index in serving sizes from the DASH trial (DASHI)[24], Alternate Mediterranean Diet Score (aMED)[25], MED Index in serving sizes from the PREDIMED trial (MEDI)[26] and Dietary Inflammation Index (DII)[27]. Similar results were also observed in the GGMP cohort for associations to single food intake frequencies (**Figure S2**). Thus, it is essential to recognize that long-term food pairing patterns serve as independent traits and may play a unique role in cardiometabolic health beyond dietary indicators.

### 3.2. Single foods exhibit limited associations with cardiometabolic traits but their pairings demonstrate pronounced effects

We then assessed the influence of long-term food pairing patterns on cardiometabolic health, including blood pressures (systolic and diastolic blood pressures, SBP and DBP) and the prevalence of hypertension, diabetes, hyperlipidemia and stroke, using 65 food intake frequencies collected from 4070 participants in the NHANES study during 2005-2010 as the discovery set (**Table S12**). Additionally, a replication set consisting of 3280 NHANES participants recruited from 1999 to 2004 was involved (**Table S12**). We used linear models to adjust for potential confounders (sex, age, race, BMI and smoking status) and further applied Spearman correlation to establish associations between long-term food pairing patterns and cardiometabolic traits. In total, 1607/1502 significant associations with 6 cardiometabolic traits at FDR < 0.05 were observed for AFPs/SFPs, comprising 58.6%/54.4% of those associations that were further replicated in the replication set (P < 0.05, **Figure 3A**).

**Figure 3.**
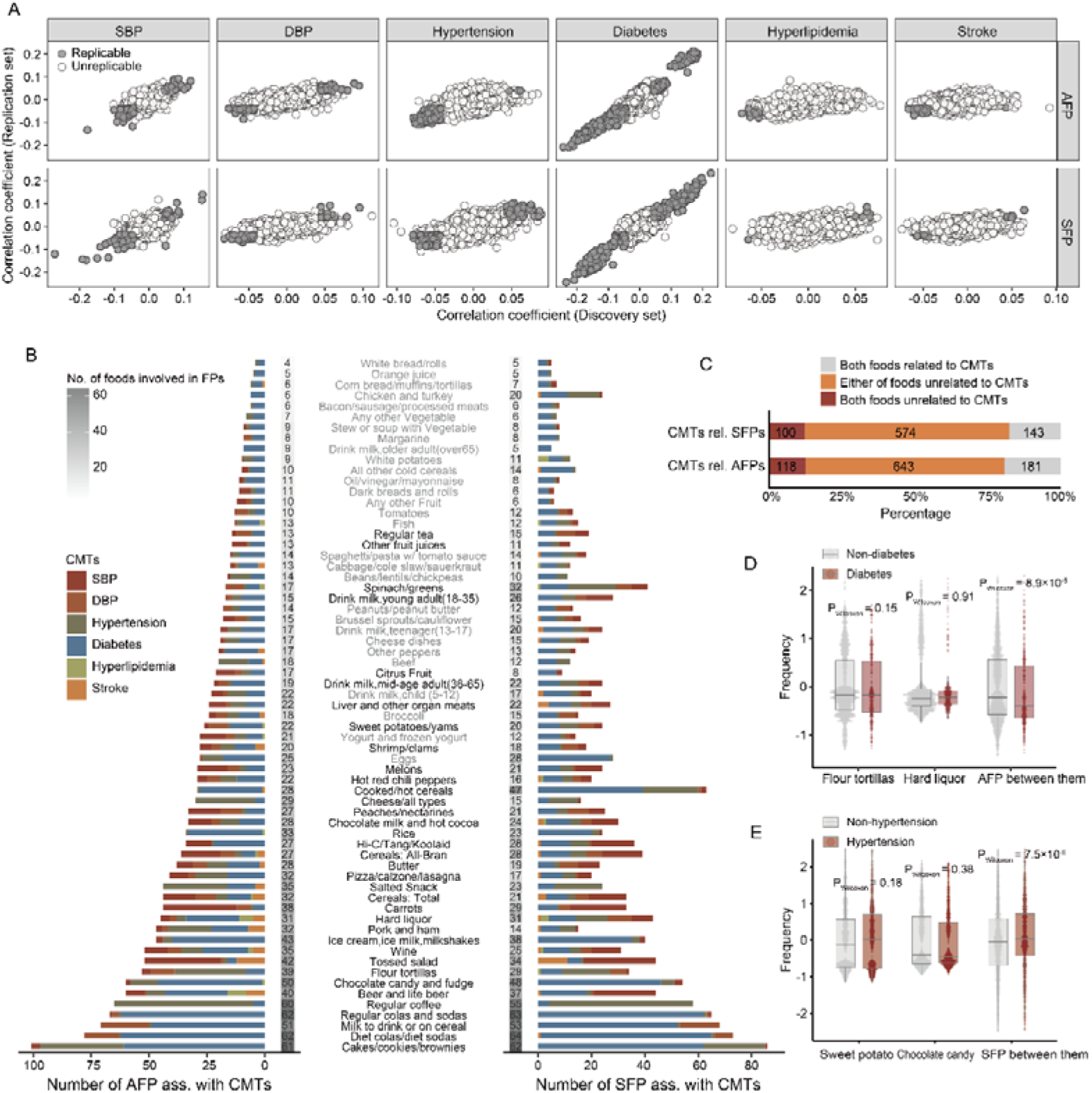
Long-term food pairing pattern associations to cardiometabolic traits. **A.** Long-term food pairing pattern associations with the 6 cardiometabolic traits in the NHANES discovery and replication sets. The X-axis indicates the Spearman correlation coefficient from the discovery set, and the Y-axis indicates the Spearman correlation coefficient from the replication set. Each dot represents one FP. Dark grey dots show replicable associations (discovery set at FDR < 0.05, replication set at P < 0.05), while light grey dots show non-replicable associations. **B.** Overview of long-term food pairing pattern associations with cardiometabolic traits per single food. The Y-axis lists all 65 different foods from food frequency questionnaires. Foods not individually associated with any of the six cardiometabolic traits are marked in grey, while those marked in black are those whose single intake frequencies are significantly associated with at least one of the six cardiometabolic traits (Spearman correlation, FDR < 0.05). The X-axis indicates the number of significant FP associations with cardiometabolic traits derived from a certain food. Numbers in the squares represent the number of single foods involved in cardiometabolic-related long-term food pairing patterns. **C.** Summary of long-term food pairing patterns with significant associations to cardiometabolic traits. Gray represents the intake frequencies of two foods within a long-term food pairing pattern both individually associated with cardiometabolic traits, while red represents interactions where neither of the two foods is individually associated with cardiometabolic traits. Orange represents pairings where either of the two foods within a long-term food pairing pattern is individually associated with cardiometabolic traits. Numbers in the bar plot represent significant additive/subtractive food pairing pattern associations to cardiometabolic traits. FP, long-term food pairing pattern; AFP, additive food pairing pattern; SFP, subtractive food pairing pattern; CMTs, cardiometabolic traits; ass., association. **D.** The additive food pairing pattern between flour tortillas and hard liquor showed a difference in diabetes prevalence, while neither the intake frequencies of the two foods showed a difference between diabetic and non-diabetic participants. The box plots show the median, first quartile, and third quartile (25th and 75th percentiles) of normalized residuals of FP levels. The upper and lower whiskers extend to the largest and smallest values no further than 1.5×IQR, respectively. Each dot represents one sample. The P value from the Wilcoxon test is shown accordingly. **E.** The subtractive food pairing pattern between sweet potato and chocolate candy showed a difference in hypertension prevalence, while neither the intake frequencies of the two foods showed a difference between hypertensive and non-hypertensive participants. The box plots show the median, first quartile, and third quartile (25th and 75th percentiles) of normalized residuals of FP levels. The upper and lower whiskers extend to the largest and smallest values no further than 1.5×IQR, respectively. Each dot represents one sample. The P value from the Wilcoxon test is shown accordingly.

In detail, the replicable associations including 143/136 SBP-related AFPs/SFPs (42 overlapped) derived from 45/49 single foods (41 overlapped), 94/75 DBP-related AFPs/SFPs (8 overlapped) derived from 39/38 single foods (32 overlapped), 210/174 hypertension-related AFPs/SFPs (74 overlapped) derived from 63/62 single foods (61 overlapped), 449/406 diabetes-related AFPs/SFPs (282 overlapped) derived from 65/65 single foods (65 overlapped), 13/7 hyperlipidemia-related AFPs/SFPs (0 overlapped) derived from 11/9 single foods (4 overlapped), and 33/19 stroke-related AFPs/SFPs (1 overlapped) derived from 16/20 single foods (7 overlapped). Notably, each single food interacted with at least one other food to form a cardiometabolic-related long-term food pairing pattern (**Figure 3B**), suggesting the broad impact of long-term food pairing patterns on human cardiometabolic health. The results of the sensitivity analysis indicated that adjustments for diet diversity had minimal impact on the findings (**Figure S3&S4**).

To ascertain whether the intake frequency of single foods is associated with those cardiometabolic traits without constructing food pairing patterns, we adjusted for the same confounding factors as mentioned above. Consequently, we identified 19 foods related to SBP, 12 to DBP, 12 to hypertension, 11 to diabetes, 1 to hyperlipidemia, and 4 to stroke (discovery set at FDR<0.05, replication set at P<0.05, **Figure 3B, Table S13**). By comparing the cardiometabolic association results with single foods and long-term food pairing patterns, we found that 80.8%/82.5% of cardiometabolic-related AFPs/SFPs were formed by single foods that were not individually associated with cardiometabolic traits (**Figure 3C**). For instance, the intake frequencies of both flour tortillas and hard liquor were not individually associated with the risk of diabetes (**Figure 3D**). However, the additive food pairing pattern (AFP) between the two foods showed a significantly negative association with the risk of diabetes (**Figure 3D**). Besides, the intake frequencies of both sweet potato and chocolate candy were not individually associated with the risk of hypertension (**Figure 3E**). However, the subtractive food pairing pattern (SFP) between the two foods showed a significantly positive association with the risk of hypertension (**Figure 3E**).

Additionally, we observed that consistent associations between single foods and cardiometabolic traits may change or even reverse when their pairing patterns are considered. For instance, higher intake frequencies of hot coca and diet cola/soda were both associated with an increased risk of diabetes (**Figure S5**). However, the SFP between them showed negative associations with the risk of diabetes(**Figure S5**). Therefore, the proposed long-term food pairing patterns provide an additional layer of information that is independent of single food intake in relation to human cardiometabolic health.

### 3.3. Long-term food pairing pattern associations to cardiometabolic traits at hyper food classification group level exhibit consistency across populations

We further examined whether the observed cardiometabolic-related long-term food pairing patterns in the westernized NHANES cohort show consistency in our GGMP cohort with eastern dietary preferences. Using the same strategy, we established 104/89 SBP-related AFPs/SFPs, 36/30 DBP-related AFPs/SFPs, 24/20 hypertension-related AFPs/SFPs, 2/1 diabetes-related AFPs/SFPs, 1 hyperlipidemia-related SFPs, and 1 stroke-related SFPs (FDR < 0.05, **Figure 4A**). By comparing the number of identified cardiometabolic-related long-term food pairing patterns, we observed that the top long-term food pairing pattern associations were mainly related to hypertension-related traits, including SBP, DBP, and the prevalence of hypertension between the two cohorts (**Figure 4A**).

**Figure 4.**
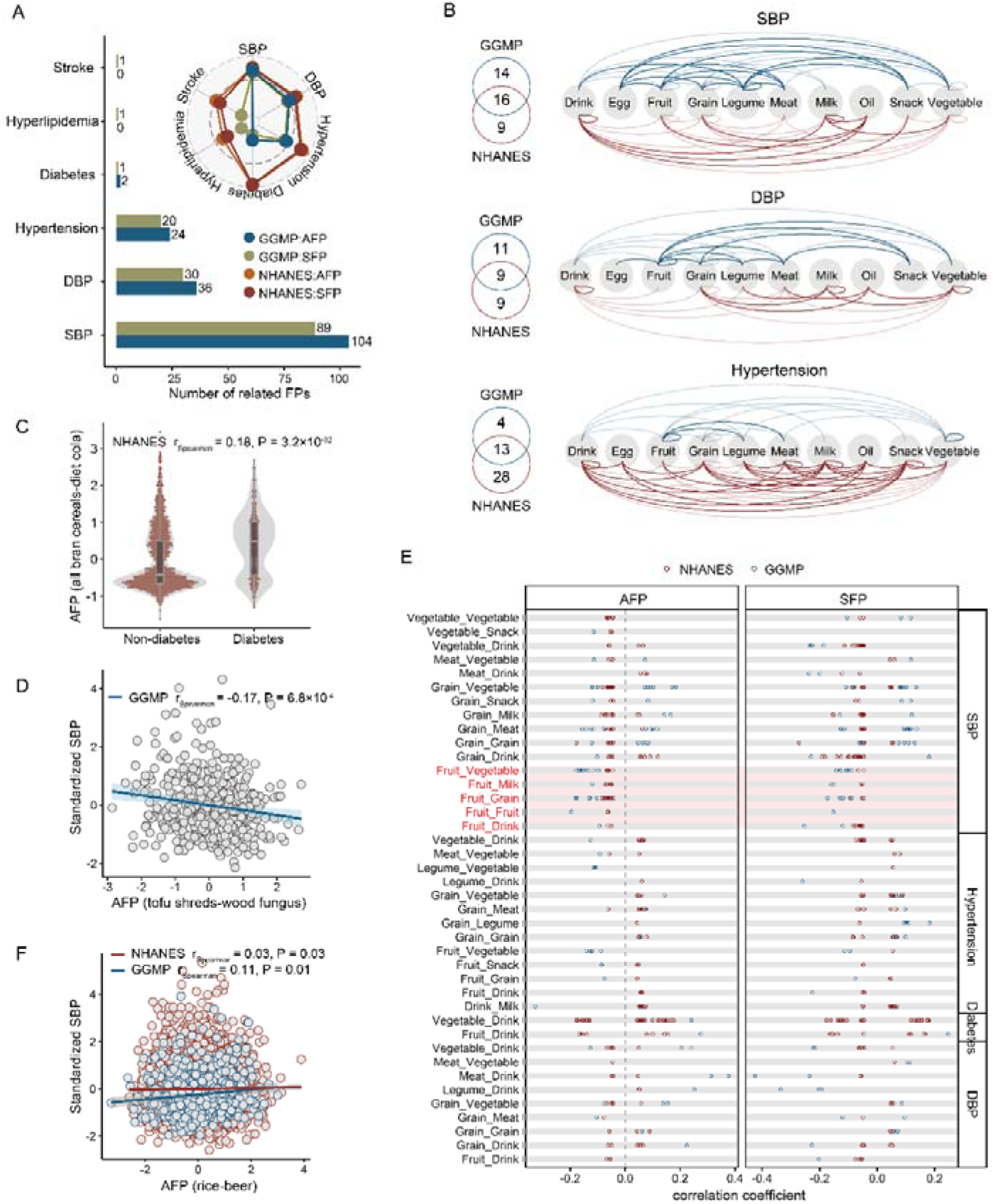
Cardiometabolic trait-related long-term food pairing patterns show consistency at hyper food classification group level between populations. **A.** Summary of long-term food pairing pattern associations with cardiometabolic traits between populations. The bar plot shows the number of long-term food pairing pattern associations with cardiometabolic traits in the GGMP cohort. The radar plot compares the number of significant long-term food pairing pattern associations with cardiometabolic traits between the NHANES and GGMP cohorts. **B.** Comparison of long-term food pairing pattern associations with cardiometabolic traits at hyper food classification group level between populations. The Venn diagram summarizes the overlap of significant long-term food pairing pattern associations with a specific cardiometabolic trait at hyper food classification group level between populations. The arc diagram on the left illustrates the detailed food pairing patterns at hyper food classification group level between populations. Darker arcs indicate non-overlapping associations, while lighter arcs represent shared associations. **C.** The additive food pairing pattern between all-bran cereals and diet colas shows a difference in diabetes prevalence. The box plots display the median, first quartile, and third quartile (25th and 75th percentiles) of normalized residuals of AFP levels. The upper and lower whiskers extend to the largest and smallest values within 1.5×IQR, respectively. Each dot represents one sample. The P value from the Spearman test is shown accordingly. **D.** The subtractive food pairing pattern between tofu shreds and wood fungus is negatively associated with systolic blood pressure. The X and Y axes indicate the normalized residuals of systolic blood pressure and the subtractive food pairing pattern between tofu shreds and wood fungus, respectively. Each dot represents one sample. The line shows the linear regression line, with gray shading representing the 95% confidence interval (CI). The Spearman correlation coefficient and P value are shown accordingly. **E.** Individual long-term food pairing pattern associations with cardiometabolic traits at hyper food classification group level between populations. Each dot represents an individual long-term food pairing pattern association with a specific cardiometabolic trait, grouped at hyper food classification level. The left Y-axis indicates long-term food pairing pattern at hyper food classification group level based on individual long-term food pairing patterns, and the right Y-axis indicates cardiometabolic traits with significant associations with long-term food pairing patterns. Long-term food pairing patterns at hyper food classification group level marked in red show the same association directions with cardiometabolic traits between populations. The X-axis indicates the Spearman correlation coefficient of long-term food pairing pattern associations with cardiometabolic traits. **F.** The subtractive food pairing pattern between rice and beer is positively associated with SBP in both populations. The X-axis indicates normalized residuals of long-term food pairing pattern, and the Y-axis indicates normalized systolic blood pressure. Each dot represents one sample. The line shows the linear regression line, with gray shading representing the 95% confidence interval (CI). The Spearman correlation coefficient and P value are shown accordingly.

Given the fact that only a fewer number of dietary habits were completely consistent between NHANES and GGMP cohorts (**Table S3&S4**), we broadened our perspective and analyzed the similarity of cardiometabolic-related long-term food pairing patterns between the two cohorts by categorizing long-term food pairing patterns into 10 hyper food groups, including milk, grain, meat, fruit, and others (**Figure 4B**). In doing so, we observed that 16, 9, 13, and 2 pairing patterns between hyper food classification group derived from long-term food pairing patterns, associated with SBP, DBP, hypertension, and diabetes, respectively, were shared in both cohorts (**Figure 4B**). Of note, we also observed many non-overlapping cardiometabolic-related long-term food pairing patterns at the hyper food classification group level (**Figure 4B**) between the two cohorts, indicating potential regional-specific long-term food pairing patterns on human cardiometabolic health. For instance, the AFP of all-bran cereals and diet colas/sodas was positively associated with the risk of diabetes in the NHANES cohort (r_Spearman_ = 0.18; P = 3.2×10^-32^) (**Figure 4C**), while the AFP of tofu shreds and wood fungus was negatively associated with SBP in the GGMP cohort (r_Spearman_ = -0.17; P = 6.8×10^-4^) (**Figure 4D**).

When plottingt he effect size of each cardiometabolic-related long-term food pairing patterns at the hyper food classification group level (**Figure 4E**), we observed that many foods from the fruit category, interacting with categories such as vegetables, drinks, grains, and milks, exhibited consistent negative associations with SBP between populations (**Figure 4E**, marked in red). Besides, we observed that despite some long-term food pairing patterns showing large variations between cohorts, they exhibited consistent associations with cardiometabolic traits. For example, the AFP of rice and beer showed substantial variation between populations (**Figure 2A**) but demonstrated consistent positive associations with SBP in both the NHANES cohort and GGMP cohort (**Figure 4F**). This underscores the consistent effects of long-term food pairing patterns on cardiometabolic traits among populations with diverse eating habits at hyper food classification group level. It provides valuable data for designing personalized dietary strategies that consider long-term food pairing patterns at hyper food classification group level to enhance cardiometabolic health.

### 3.4. The impact of long-term food pairing patterns on gut microbiome composition

We recently demonstrated that the gut microbiome significantly contributes to host cardiac metabolism, as evidenced by its impact on inter-individual variations in various categories of plasma metabolites[9,34]. Thus, characterizing the influence of long-term food pairing patterns on the composition of the gut microbiome may provide mechanistic insights into the etiologies of human cardiometabolic disorders. By associating long-term food pairing patterns with 205 microbial genera with a prevalence rate of more than 10% in the GGMP cohort (n=6,994), while adjusting for gender, age, BMI, and smoking status, we identified 3575 associations between 734 AFPs and 179 genera, and 2728 associations between 585 SFPs and 196 genera, respectively (FDR<0.05, **Table S7**). In the meanwhile, we also examined the association between single foods and gut microbiomes without constructing long-term food pairing patterns. In total, we identified 854 associations between 196 gut microbial genera and 56 foods (FDR<0.05, **Table S8**).

Interestingly, we observed that long-term food pairing patterns have broad associations with genera from *Clostridiales*, *Burkholderiales*, *Bacteroidales*, and the *Erysipelotrichales* order (**Figure 5A**), as reflected by distinct clusters in the association heatmaps (**Figure S6&S7**). However, their associations with single foods were rather sparse (**Figure S6&S7**), indicating that the gut microbiome may have a propensity for specific long-term food pairing patterns independent of individual foods. For instance, microbial associations with long-term food pairing patterns derived from fresh fruit encompassed not only all 33 fresh fruit-related genera but also numerous genera unrelated to fresh fruit (**Figure 5B**). Furthermore, specific foods paired with others to be widely associated with gut microbiome composition, but their individual associations with microbial genera were limited (**Figure 5C**). For example, long-term food pairing patterns formed by mutton and 26 other foods were linked to 45 genera, however, mutton itself was not directly linked to any genus (**Figure 5C**). Additionally, the number of genera associated with long-term food pairing patterns derived from tofu exceeded eight times the number of genera specifically related to tofu (**Figure 5C**). Moreover, the proposed AFPs and SFPs also have independent associations with gut microbial composition, as reflected by the non-overlapping associations (**Figure 5D**), indicating that these proposed two types of long-term food pairing patterns have unique effects on the gut microbiome composition. For instance, AFP between rice and other cereals was linked to 36 different genera, which were not associated with their SFP. Conversely, the SFP between rice and other fried foods was linked to 42 different genera but not associated with their AFP (**Figure 5D**).

**Figure 5.**
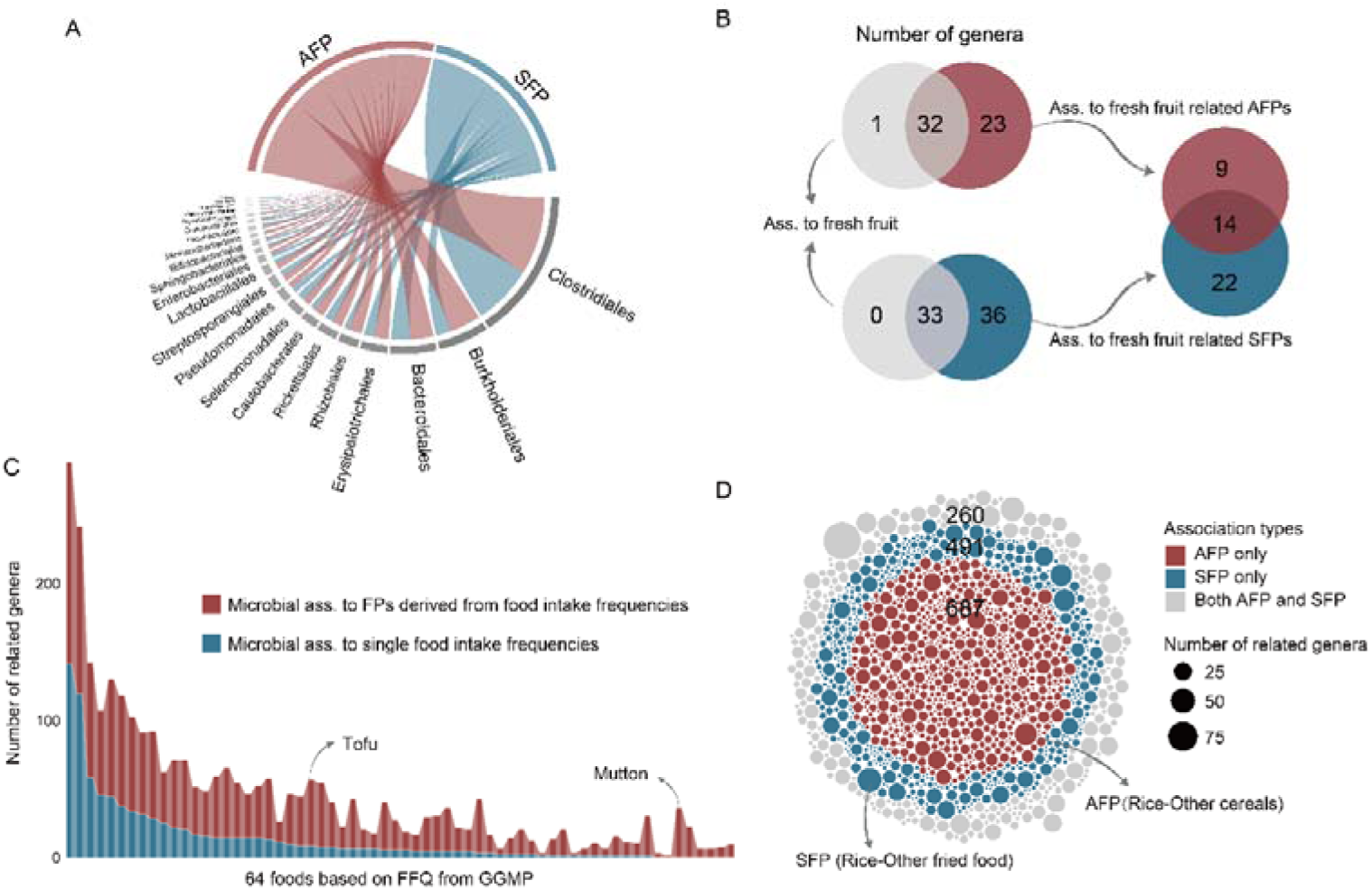
The influence of long-term food pairing patterns on gut microbiome composition. **A.** Summary of long-term food pairing pattern associations with the composition of gut microbial genera. The associated additive food pairing patterns are colored red, and the associated subtractive food pairing patterns are colored blue. Each line indicates a significant association between a food pairing pattern and gut microbial genera within the same order (Spearman correlation, FDR < 0.05). **B.** Comparison of microbial genera associated with fresh fruit intake frequency and long-term food pairing patterns derived from fresh fruit. The Venn diagram on the left indicates the overlap of microbial genera associated with fresh fruit intake frequency and long-term food pairing patterns derived from fresh fruit. The diagram on the right shows the overlap of microbial genera associated only with additive or subtractive food pairing patterns **C.** Comparison of microbial associations with intake frequency of single foods and long-term food pairing patterns derived per single food. The X-axis lists all 64 different foods from food frequency questionnaires in the GGMP cohort. The Y-axis indicates the number of microbial genera with significant associations with single food intake frequencies and food pairing patterns derived per single food. **D.** Overlap of long-term food pairing pattern associations with microbial genera between additive and subtractive food pairing patterns. Microbial associations with both additive and subtractive food pairing patterns are marked in grey, while those only associated with additive or subtractive food pairing patterns are marked in red and blue, respectively. Each dot represents one food pairing pattern, with the dot size indicating the number of related genera.

### 3.5. The gut microbiome mediates associations between long-term food pairing patterns and cardiometabolic traits

Of the 220 unique long-term food pairing patterns associated with cardiometabolic traits and the 1319 associated with microbes, 192 food pairing patterns are associated with both (**Figure 6A**). To evaluate whether microbes can mediate the long-term food pairing patterns impact on cardiometabolic traits, we associated 205 gut microbial genera with the 6 cardiometabolic traits and established 668 significant associations with 190 genera (FDR < 0.05, **Figure 6B**, **Table S9**), including 138 SBP-related genera, 116 DBP-related genera, 127 hypertension-related genera, 113 diabetes-related genera, 126 hyperlipidemia-related genera and 48 stroke-related genera.

**Figure 6.**
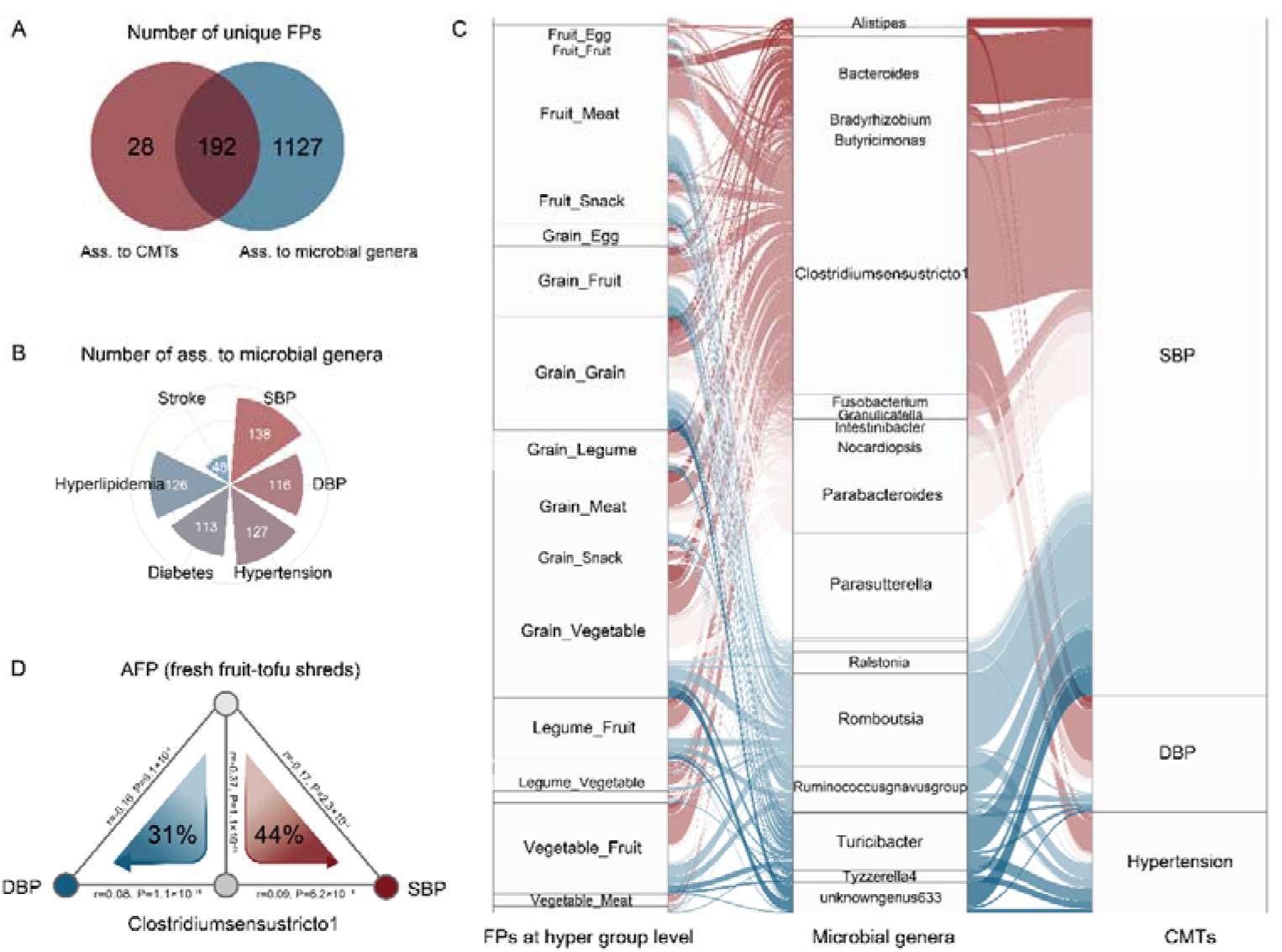
The gut microbiome mediates associations between long-term food pairing patterns and cardiometabolic traits. **A.** Overlap of long-term food pairing patterns with significant associations to cardiometabolic traits and gut microbial genera. **B.** Summary of microbial associations with different cardiometabolic traits. The number indicates significant associations at FDR < 0.05. **C.** Gut microbial genera mediate long-term food pairing pattern associations with cardiometabolic traits. The sankey plot illustrates significant mediation effects of the gut microbiome at FDR < 0.05. Shown are long-term food pairing patterns at hyper food classification group level (left), microbial genera (middle), and cardiometabolic traits (right). Curved lines connecting the panels denote significant mediation linkages, with colors representing different gut microbes acting as mediators. **D.** *Clostridiumsensustricto1* mediates the additive food pairing pattern between fresh fruit and tofu shreds in association with both systolic and diastolic blood pressures. The gray lines indicate the associations between the two factors, with corresponding Spearman coefficients and P values. The mediation effects are shown with arrows.

We then employed mediation analysis to explore the pathways through which long-term food pairing patterns influence host cardiometabolic traits via gut microbe. Our analysis identified 755 mediation linkages, with 160 long-term food pairing pattern features (comprising 86 AFPs and 74 SFPs) significantly affecting host cardiometabolic traits through 31 gut microbial genera (FDR < 0.05, **Figure 6C**). Notably, we found that out of the 220 unique long-term food pairing patterns linked to cardiometabolic traits, 72.7% demonstrated a noteworthy influence on these traits via the gut microbiome (**Table S10**). This underscores the critical role of the gut microbiome in mediating the impact of long-term food pairing patterns on cardiometabolic health.

Besides, a substantial portion (26.6%) of these mediated linkages were associated with long-term food pairing pattern impact on blood pressure via the *Clostridiumsensustricto1* genus. Clinical studies have implicated *Clostridiumsensustricto1* in amino acid and central carbohydrate metabolism[35], as well as its potential to raise host blood pressure[36]. Here, we observed that the AFP between fresh fruit and tofu shreds contributed to a decrease in SBP and DBP by reducing the abundance of *Clostridiumsensustricto1* (SBP: 44%, P_mediation_ < 0.001; DBP: 31%, P_mediation_ <0.001; **Figure 6D**). Our findings thus suggest that *Clostridiumsensustricto1* may play a role in mediating the effect of long-term food pairing patterns on cardiometabolic health.

### 3.6. The mediation effects of gut microbiome rely on metabolic pathways

To delve into the functional role of gut microbes in mediating the impact of long-term food pairing patterns on cardiometabolic traits, we examined how these microbes influence such traits through their metabolic pathways. Among the 31 genera identified as mediators of long-term food pairing patterns on cardiometabolic traits (FDR < 0.05, **Table S10**), we characterized 237 metabolic pathways associated with both cardiometabolic traits and microbial genera with mediator roles (FDR < 0.05, **Table S11**).

Further utilizing mediation analysis, we identified 4019 significant mediation linkages involving 212 pathways that mediate the impact of microbes on cardiometabolic traits (FDR < 0.05, **Table S11**). Notably, we observed that microbes mediate the impact of long-term food pairing patterns on cardiometabolic traits through pathways involved in well-known short-chain fatty acid and amino acid biosynthesis. For instance, propionic acid, a short-chain fatty acid produced from dietary fiber by the gut microbiome, is known to play a crucial role in cardiometabolic health[14,37]. Our results indicated that the genera *Alistipes*, *Butyricimonas*, and *Parasutterella* contributed to mediating the impact of long-term food pairing patterns derived from fiber-enriched vegetables and fruits on systolic blood pressure through the pyruvate fermentation to propanoate I pathways (**Figure 7A**, **Figure 7C**).

**Figure 7.**
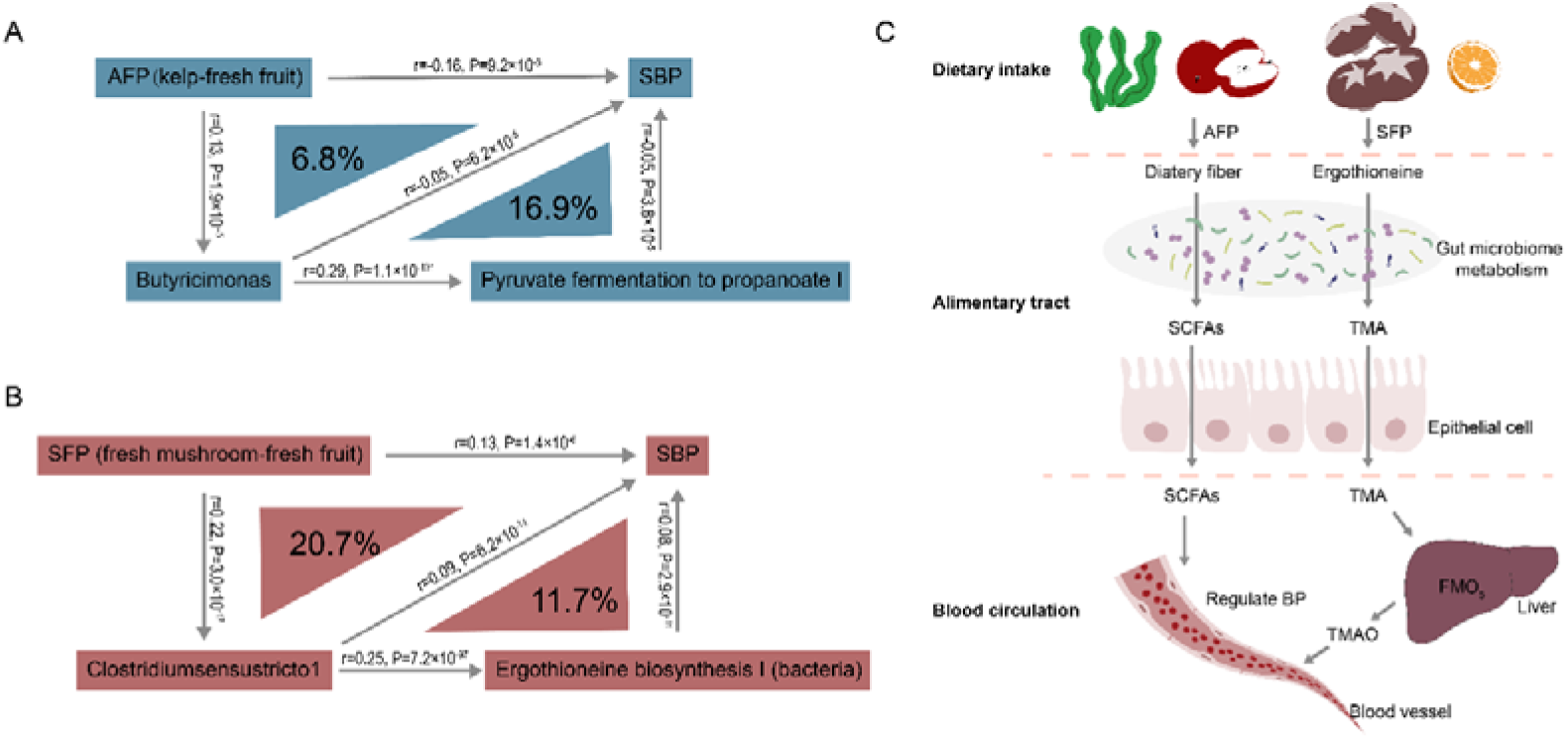
The medication effects of gut microbiome in long-term food pairing pattern associations to cardiometabolic traits rely on microbial metabolic pathways. **A.** The additive food pairing pattern between kelp and fresh fruit is associated with systolic blood pressure through propanoate biosynthesis by *Butyricimonas*. The gray lines indicate the associations between the two factors, with corresponding Spearman coefficients and P values. The mediation effects of specific gut microbiome and its metabolic pathway are shown with arrows. **B.** The subtractive food pairing pattern between fresh mushrooms and fresh fruit is associated with systolic blood pressure through ergothioneine biosynthesis by *Clostridiumsensustricto1*. The gray lines indicate the associations between the two factors, with corresponding Spearman coefficients and P values. The mediation effects of specific gut microbiome and its metabolic pathway are shown with arrows. **C.** Graphical illustration of the two examples mentioned above.

Additionally, previous studies have shown that gut microbiomes may promote the synthesis of trimethylamine (TMA) by utilizing ergothioneine from the host’s daily diets[38], potentially leading to elevated blood pressure[39]. Here, we observed that 22 distinct gut microbial genera mediated the impact of long-term food pairing patterns on cardiometabolic traits through the ergothioneine biosynthesis I pathway (**Table S11**). Human beings primarily obtain ergothioneine from mushrooms[40]. Interestingly, we found that long-term food pairing patterns derived from ergothioneine-rich mushrooms influenced host blood pressure by affecting certain potentially pathogenic microbiomes capable of absorbing and metabolizing ergothioneine. Specifically, a higher host SFP of fresh mushroom sand fresh fruits (intaking more mushrooms and less fruits) elevated host systolic blood pressure by increasing the abundance of certain potentially pathogenic genera belonging to the *Clostridiales* order, including *Clostridiumsensustricto1*, *Romboutsia*, and *Tyzzerella4* (**Table S10**). These genera acted through the ergothioneine biosynthesis I pathway to enhance their colonization and TMA synthesis, thereby contributing to the observed increase in host blood pressure (**Figure 7B**, **Figure 7C**, **Table S11**). Taken together, these findings provide valuable insights into the functional role of gut microbes in mediating the impact of long-term food pairing patterns on cardiometabolic traits through metabolic pathways.

## 4. Discussion

It is widely acknowledged that both the consumption of certain foods such as red meat[4,41] and dietary patterns like DASH (Dietary Approaches to Stop Hypertension)[6] and MED (Mediterranean Diet)[7] are associated with cardiometabolic health. In contrast to previous dietary indicators, the concept of long-term food pairing pattern proposed here aims to elucidate the effects of balanced and imbalanced combinations of different foods on human cardiometabolic phenotypes, including additive food pairing pattern and subtractive food pairing pattern. Individual long-term food pairing pattern shows weak correlation with classical dietary indicators such as dietary intake frequency or dietary pattern indices, it operates independently of these indicators in principle.

Using two large-scale population-based cohorts, we investigated associations between long-term food pairing patterns and cardiometabolic traits, observing consistent results across populations with vastly different dietary preferences. Specifically, we identified 942/817 and 166/142 cardiometabolic-related AFPs/SFPs in the NHANES cohort and our GGMP cohort after adjusting for potential confounding factors. Importantly, the consistency of pairings between hyper food classification groups, to which the aforementioned cardiometabolic-related long-term food pairing patterns belong, was evident across two cohorts characterized by entirely disparate dietary habits. For instance, both cohorts supported the negative association between the fruit category and drinks, grains, milks, and vegetables with SBP, indicating that increased pairing of fruits with these food groups (higher AFPs) benefits host blood pressure. Moreover, the effects are enhanced when the consumption frequency of the latter exceeds that of fruits (higher SFPs). These findings underscore the complexity and similarity of real-world long-term food pairing patterns, emphasizing the importance of considering the intricate inter play among daily diets in understanding their impact on human cardiometabolic health.

Importantly, our observations highlight that individual foods forming long-term food pairing patterns consistently engage with other diets linked to cardiometabolic traits, contrasting with a lower occurrence when considering individual foods alone. Additionally, correlations between foods and cardiometabolic traits may change or reverse upon constructing long-term food pairing patterns. For instance, the positive association between diabetes and hot cocoa, as well as diet cola/soda, can reverse when these foods participate in constructing SFP. This underscores the potential oversight of prior studies focusing solely on single food-health relationships, neglecting the broader impact of diet-to-diet pairings. Further investigation is needed to determine if these findings can inform nutritional guidancei n dietary practices, potentially through targeted interventions, longitudinal studies, or controlled trials assessing the practical implications of observed long-term food pairing patterns on cardiometabolic health.

To gain mechanistic insights into the impact of long-term food pairing patterns on human cardiometabolic traits, we examined the mediating role of gut microbes and their metabolic pathways. We found that certain foods paired with others, exhibiting broad associations with microbial genera, surpassing their individual associations with gut microbiomes. This highlights the comprehensive nature of long-term food pairing patterns in relation to diet-associated gut microbiomes. Our mediation analysis identified 755 linkages, with 160 long-term food pairing patterns affecting host cardiometabolic traits through 31 gut microbiomes, indicating the moderating role of gut microbiome-mediated long-term food pairing patterns on host blood pressure. Furthermore, analysis of gut microbial metabolic pathways revealed that *Alistipes*, *Butyricimonas*, and *Parasutterella* may mediate the effects of long-term food pairing patterns from fiber-enriched vegetables and fruits on systolic blood pressure, possibly through pyruvate fermentation to propanoate I pathways[14,37]. Additionally, long-term food pairing patterns involving ergothioneine-rich mushrooms[40] influenced host blood pressure by metabolizing ergothioneine to trimethylamine biosynthesis[38], leading to increased blood pressure[39]. These findings elucidate potential mechanisms underlying the impact of long-term food pairing patterns on human cardiometabolic health.

We acknowledge several limitations in our study. Firstly, our study employs an observational design, inherently susceptible to unknown factors, despite adjusting for existing confounding variables to minimize interference. Independent replications in other cohorts with diverse settings could further bolster the robustness of the conclusions[42–44]. Secondly, long-term food pairing pattern calculations rely on food frequency questionnaires (FFQs), which have inherent limitations. FFQs are subject to recall bias and measurement error, potentially leading to misclassification of food frequencies. A common issue is under-reporting, where participants may underestimate their intake due to social desirability bias or memory limitations. Finally, while we have demonstrated a link between gut microbiota and associations among long-term food pairing patterns and cardiometabolic health, the underlying causal relationship remains unclear. Despite these limitations, our study provides a novel perspective on the potential role of long-term food pairing patterns in human cardiometabolic health and its underlying microbial insights. The long-term food pairing patterns reported here offer a comprehensive resource that can guide follow-up studies aimed at designing preventive and therapeutic precision nutritional strategies for human cardiometabolic health.

## 5. Conclusions

Our data suggest that balance between long-term food pairing patterns is broadly associated with cardiometabolic traits by modulating gut microbial functionalities, in contrast to the lower occurrence when considering individual foods in isolation. These findings offer a novel perspective for designing personalized dietary strategies that extend beyond current dietary indicators to enhance cardiometabolic health.

## Acknowledgements

We thank the participants and staff of the GGMP and NHANES for their collaboration. This project was funded by the National Natural Science Foundation of China (NSFC, 32394052, 32270077 and Excellent Young Scientists Fund Program Overseas-2022); Jiangsu Shuang chuang Project (Medical Innovation Team, Medical Expert & JSSCBS20221815); the Natural Science Foundation of Jiangsu(BK20220709); the Nanjing Medical University (303073572NC21, YNRCZN0301, CMCM202204 and GSKY20210105); Suzhou Science and Technology Development Plan Foundation (SKY2021010); Gusu Health Talent Program (GSWS202206); Development of Jiangsu Higher Education Institutions Priority Academic Program (PAPD). The funders had no role in the study design, data collection and analysis, decision to publish, or preparation of the manuscript.

## Compliance with ethics guidelines

We thank the participants and staff of the GGMP and NHANES for their collaboration. H.Z., X.K., and L.C. conceptualized and managed the study. W.W., Q.D., J.F., F.X., Q.D. and A.L. generated the data. Q.D., Y.S., J.F., F.X. and L.C. analyzed the data. Q.D., Y.S., M.G. and L.C. drafted the manuscript. Q.D., Y.S., M.G., J.F., F.X., Y.J., Y.Z., B.M., L.L., X.W., Q.D., T.W., H.Z., J.S., Y.W., Y.S., L.T., W.S., A.L., S.H., J.Z., Y.H., H.Z., W.W., X.K. and L.C. reviewed and edited the manuscript.

## Data and code availability

The National Health and Nutrition Examination Survey (NHANES) datasets are available in the NHANES repository (https://wwwn.cdc.gov/nchs/nhanes/). The gut microbiome sequencing data of the Guangdong Gut Microbiome Project is deposited at the European Nucleotide Archive (https://www.ebi.ac.uk/ena/) with accession number PRJEB18535. The analysis codes are available via: https://github.com/MicrobiomeCardioMetaLab/FoodInteractions_project.

## Supplemental Materials

Data Supplement Figures 1-7

Data Supplement Tables 1-13

## Supplementary figures

**Figure S1.**
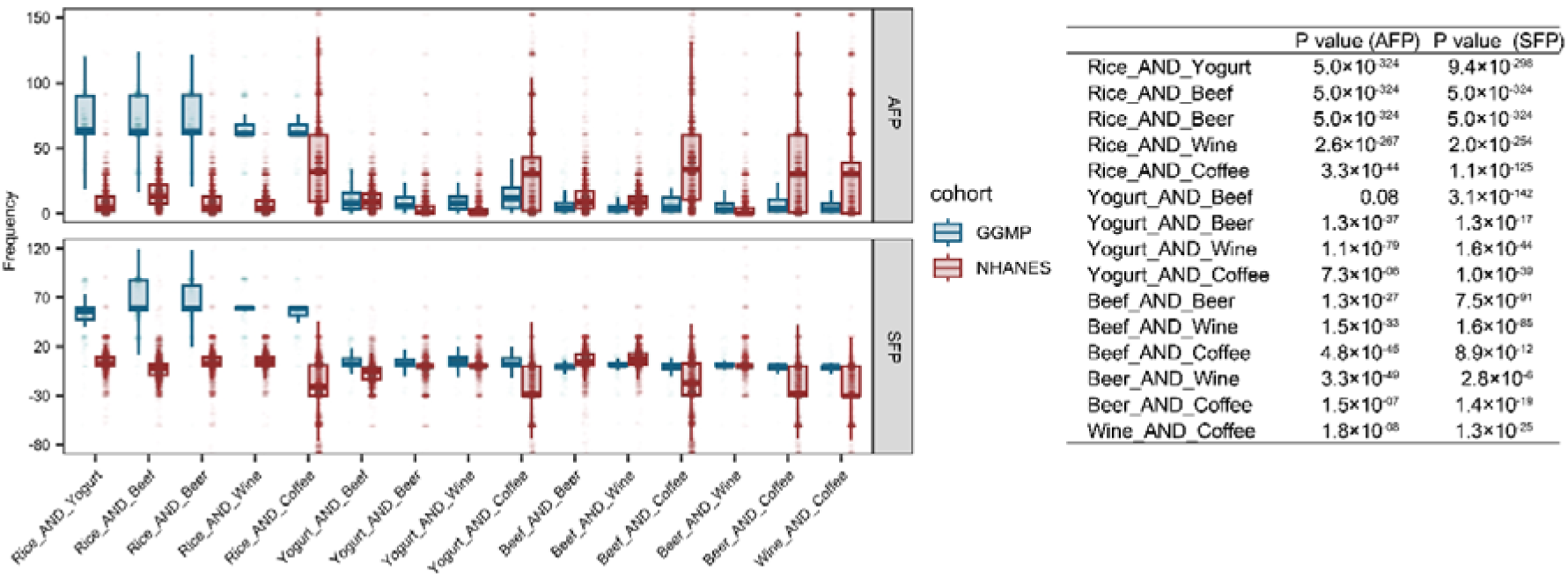
Distribution of overlapped long-term food pairing patterns between populations. The boxplot displays the median, first quartile, and third quartile (25th and 75th percentiles) of the food pairing pattern levels. The upper and lower whiskers extend to the largest and smallest values no further than 1.5× the IQR, respectively. Each dot represents one sample. The P values from the Wilcoxon test are shown on the right side.

**Figure S2.**
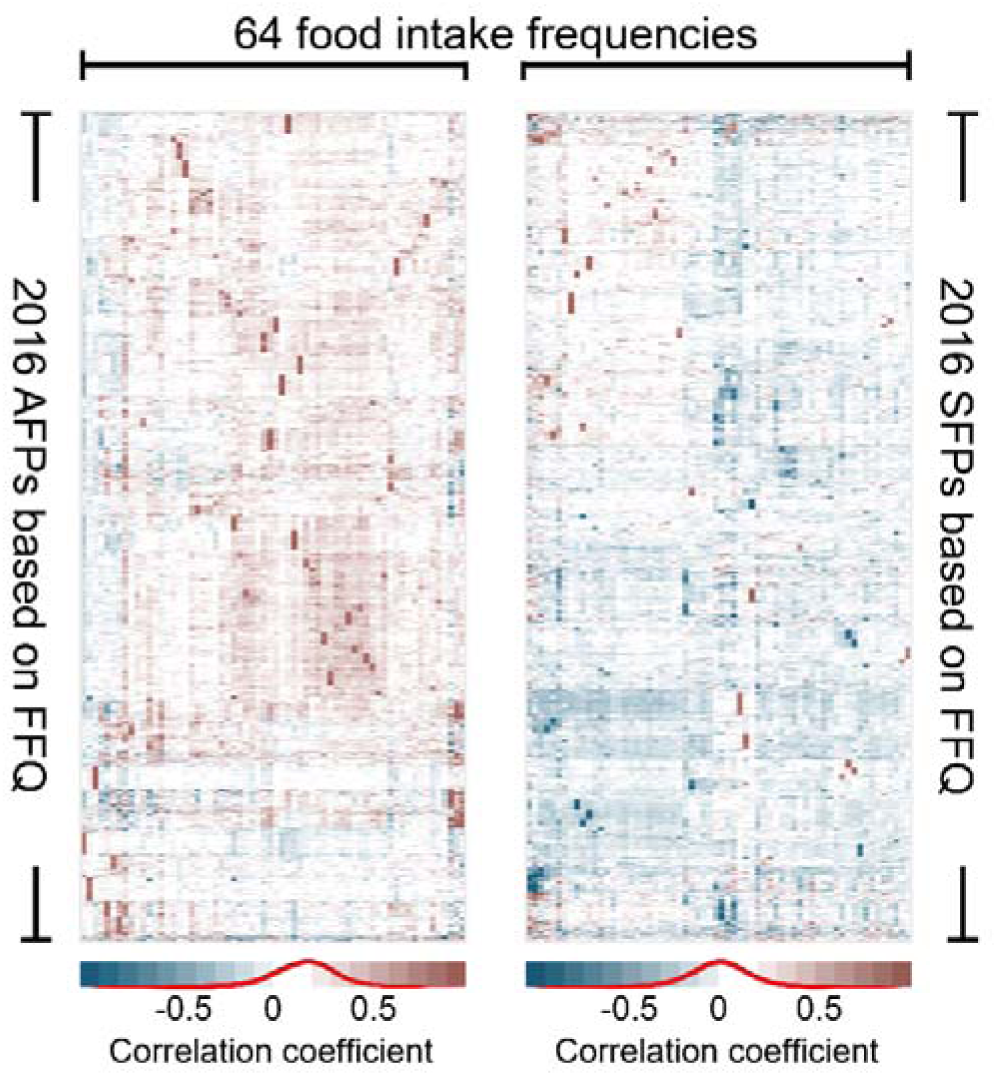
Heatmap illustrating Spearman correlations between 64 single food intake frequencies and 2016 long-term food pairing patterns in the GGMP. The intensity of color represents the correlation strength.

**Figure S3.**
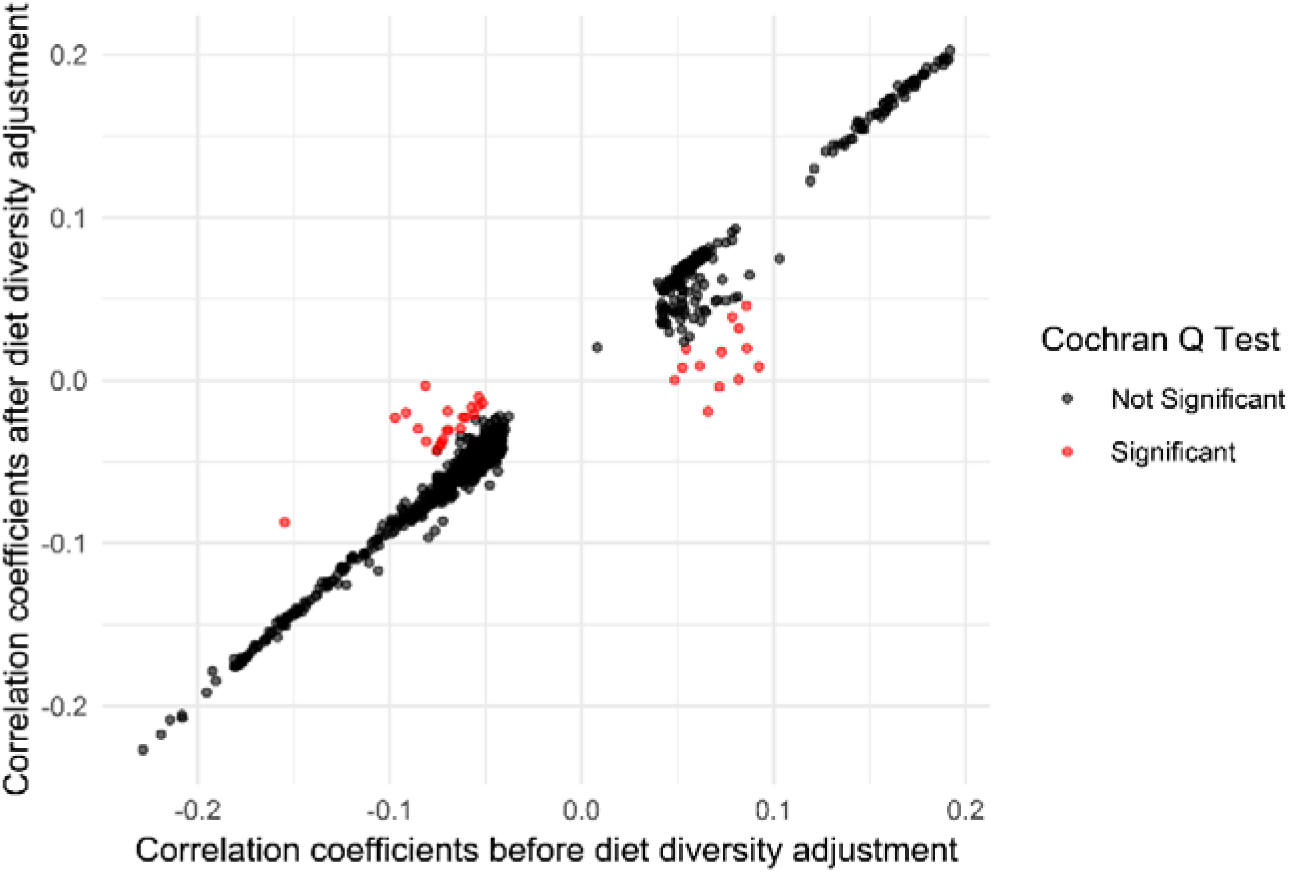
The scatter plot illustrates the comparison of correlation coefficients between additive food pairing patterns and cardiometabolic traits before and after adjusting for dietary diversity. The x-axis represents correlation coefficients before adjustment, while the y-axis represents those after adjustment. Red dots indicate points with significant heterogeneity based on the Cochran Q test (P< 0.05), whereas black dots denote non-significant results.

**Figure S4.**
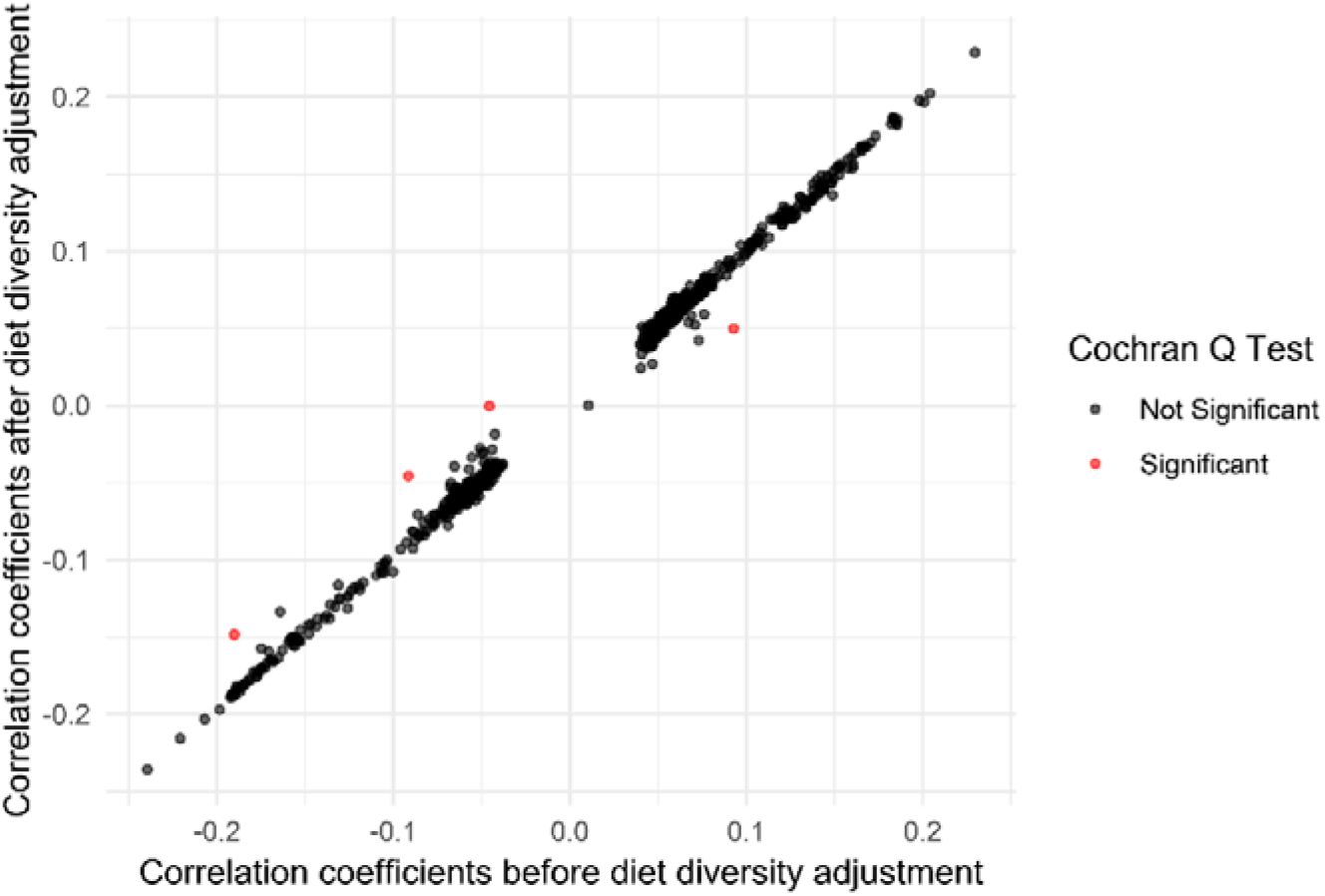
The scatter plot illustrates the comparison of correlation coefficients between subtractive food pairing patterns and cardiometabolic traits before and after adjusting for dietary diversity. The x-axis represents correlation coefficients before adjustment, while the y-axis represents those after adjustment. Red dots indicate points with significant heterogeneity based on the Cochran Q test (P< 0.05), whereas black dots denote non-significant results.

**Figure S5.**
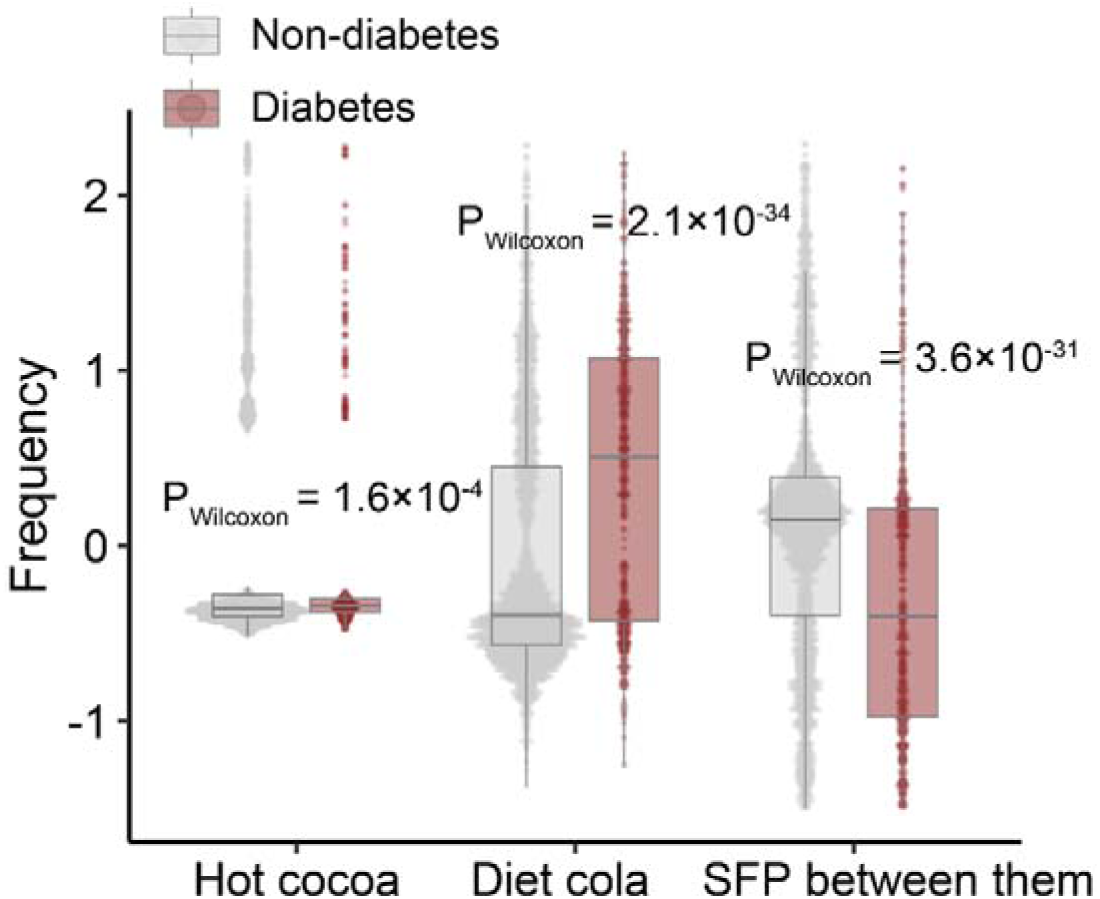
Hot cocoa and diet cola are both positively associated with the risk of diabetes, while the subtractive food pairing pattern between them is negatively associated with the risk of diabetes. The P values from the Wilcoxon test are shown accordingly.

**Figure S6.**
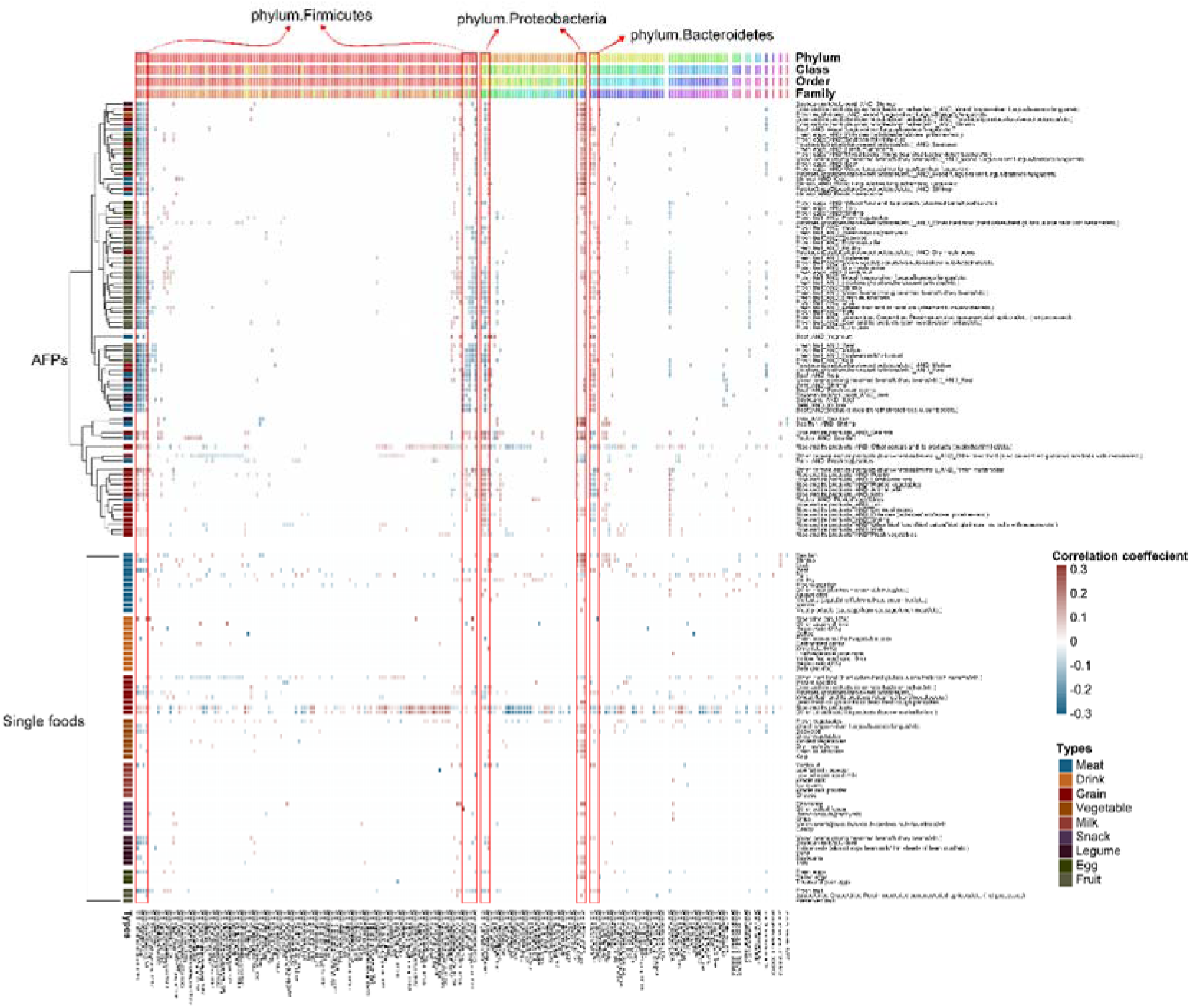
Heatmap showing microbial associations with additive food pairing patterns and their corresponding single food intake frequencies, estimated using Spearman correlation. The intensity of colors represents the correlation strength.

**Figure S7.**
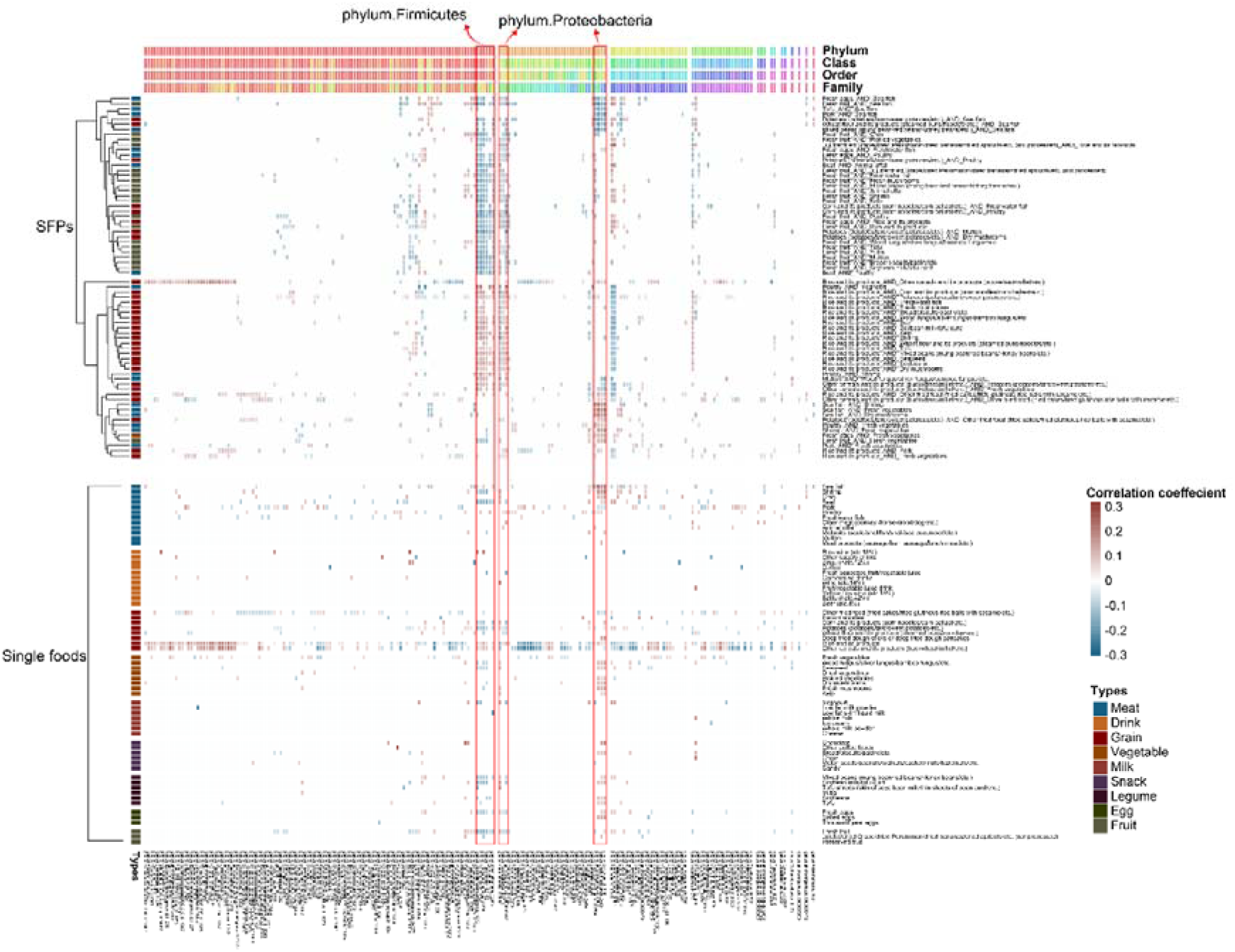
Heatmap showing microbial associations with subtractive food pairing patterns and their corresponding single food intake frequencies, estimated using Spearman correlation. The intensity of colors represents the correlation strength.

## Supplemental table legend

**Table S1.** Metadata information in the NHANES cohort.

**Table S2.** Metadata information in the GGMP cohort.

**Table S3.** Intake frequency of 65 foods in the NHANES cohort.

**Table S4.** Intake frequency of 64 foods in the GGMP cohort.

**Table S5.** Correlation between cardiometabolic traits and long-term food pairing patterns in the NHANES cohort.

**Table S6.** Correlation between cardiometabolic traits and long-term food pairing patterns in the GGMP cohort.

**Table S7.** Correlation between gut microbial genera and long-term food pairing patterns in the GGMP cohort.

**Table S8.** Correlation between gut microbial genera and single foods in the GGMP cohort.

**Table S9.** Correlation between gut microbial genera and cardiometabolic traits in the GGMP cohort.

**Table S10.** Mediation effect of gut microbiomes between long-term food pairing patterns and cardiometabolic traits in the GGMP cohort.

**Table S11.** Mediation effect of metabolic pathways between gut microbiomes and cardiometabolic traits in the GGMP cohort.

**Table S12.** Characteristics of study participants in the NHANES and GGMP cohorts.

**Table S13.** Correlation between single foods and cardiometabolic traits in the NHANES cohort.

